# Human Brain Ancestral Barcodes

**DOI:** 10.1101/2024.07.14.603450

**Authors:** Darryl Shibata

## Abstract

Dynamic CpG methylation “barcodes” were read from 15,000 to 21,000 single cells from three human male brains. To overcome sparse sequencing coverage, the barcode had ∼31,000 rapidly fluctuating X-chromosome CpG sites (fCpGs), with at least 500 covered sites per cell and at least 30 common sites between cell pairs (average of ∼48). Barcodes appear to start methylated and record mitotic ages because excitatory neurons and glial cells that emerge later in development were less methylated. Barcodes are different between most cells, with average pairwise differences (PWDs) of ∼0.5 between cells. About 10 cell pairs per million were more closely related with PWDs < 0.05. Barcodes appear to record ancestry and reconstruct trees where more related cells had similar phenotypes, albeit some pairs had phenotypic differences. Inhibitory neurons showed more evidence of tangential migration than excitatory neurons, with related cells in different cortical regions. fCpG barcodes become polymorphic during development and can distinguish between thousands of human cells.

## Introduction

Cell lineages outline tissue development. Complete fate maps are possible by direct observation for small organisms such as C. elegans, but various elegant experimental fate markers are employed for larger tissues and longer time intervals (1). For human tissues, prior experimental manipulations are impractical, and genomic alterations are employed. Somatic mutations mark subclones and their fates can be reconstructed with DNA sequencing. Recent advances in single cell technologies potentially allow fate map reconstruction at single cell resolution.

Here we show how fluctuating CpG (fCpG) DNA methylation (2) could be used as dynamic barcodes to study human brain development using single cell epigenomes annotated with their locations and phenotypes (3). DNA methylation patterns are usually copied between cell divisions, but replication errors are much higher compared to base replication, allowing for more differences between daughter cells. DNA methylation modulates expression and their patterns can be used to infer cell phenotypes (3,4), but most fCpG sites are present outside of genes or in unexpressed genes. Criteria for our fCpG barcode are as follows: 1) a defined initial pattern in a progenitor cell; 2) polymorphic changes upon cell division; 3) adequate polymorphism to distinguish between most cells; and 4) capability to record ancestry.

The brain has several features that facilitate barcode development and validation. Foremost, there is extensive single cell methylation data, with thousands of cells annotated by locations and phenotypes (3). Although billions of cells are present in an adult brain, lineage trees are compact because growth is largely neonatal. The brain also allows for serial “stopwatch” barcode sampling because development roughly follows a caudal to rostral pattern, and groups of neurons characteristically stop dividing and differentiate at different times and locations (5). Brainstem neurons emerge early (6) and their barcodes should most resemble the initial progenitor state, whereas the stopwatch runs longer for excitatory neurons that appear later in development. To facilitate presentation, barcode performance is summarized as follows: The brain fCpG barcode initializes as predominately methylated in the progenitor cell and becomes polymorphic with more diverse barcodes in excitatory neurons that emerge later in development. The barcode becomes sufficiently polymorphic to uniquely distinguish between most sampled brain cells, and barcoded cells organize into lineage trees.

## Results

### fCpG barcode identification

Barcode development was limited by the sparse single cell data (3), with < 5% of CpG sites sequenced, often with only a single read. Sparse coverage was mitigated with X-chromosome fCpG sites because only a single read can infer a binary (0,1) state in male individuals. Autosomal CpG sites require at least 2 reads to infer three possible states (0, 0.5, 1). The X-chromosome also simplifies the identification of polymorphic fCpGs because many neurons have different binary states if average methylation is between 0.25 and 0.75 in bulk WGBS adult male neurons reference data (4).

CpG sites (N∼116,000), with average methylation between 0.25 and 0.75 in bulk neurons from seven males (4), were further filtered by discarding more stable CpG sites with average methylation less than 0.2 or more than 0.8 for all cells, inhibitory neurons, and excitatory neurons in brain H02. The ∼79,000 CpGs were further filtered to remove sites with average methylation less than 0.3 or greater than 0.7 in brain H01, and ∼31,000 fCpG sites were used for analysis.

fCpG site methylation appears neutral because they are predominately intergenic, with 16% within genes or promoters (File S1). Epigenomes from 15,434 to 21,836 cells were downloaded from three male brains with a general criterion of allc.tsv.gz file sizes 90 mb or larger (Table 1). Neurons were preferentially sampled whereas glial cells were sometimes excluded (File S2). Analyzed cells had at least 500 fCpGs (average ∼1,100), with pairwise distances (PWDs) calculated between cell pairs when at least 30 fCpGs were comparable (average ∼48 fCpGs per cell pair). Each cell, annotated by its provided phenotype and location, is characterized by its fCpG methylation level and its PWDs from other cells. A PWD of 0 is a perfect match and 0.5 indicates randomization. fCpG methylation was variable between cells with averages of ∼58% for all three brains (Fig 1A). The 73 to 197 million possible cell pair comparisons revealed polymorphic barcodes with average PWDs of ∼0.47 between cells (Fig 1B).

**Fig 1:**
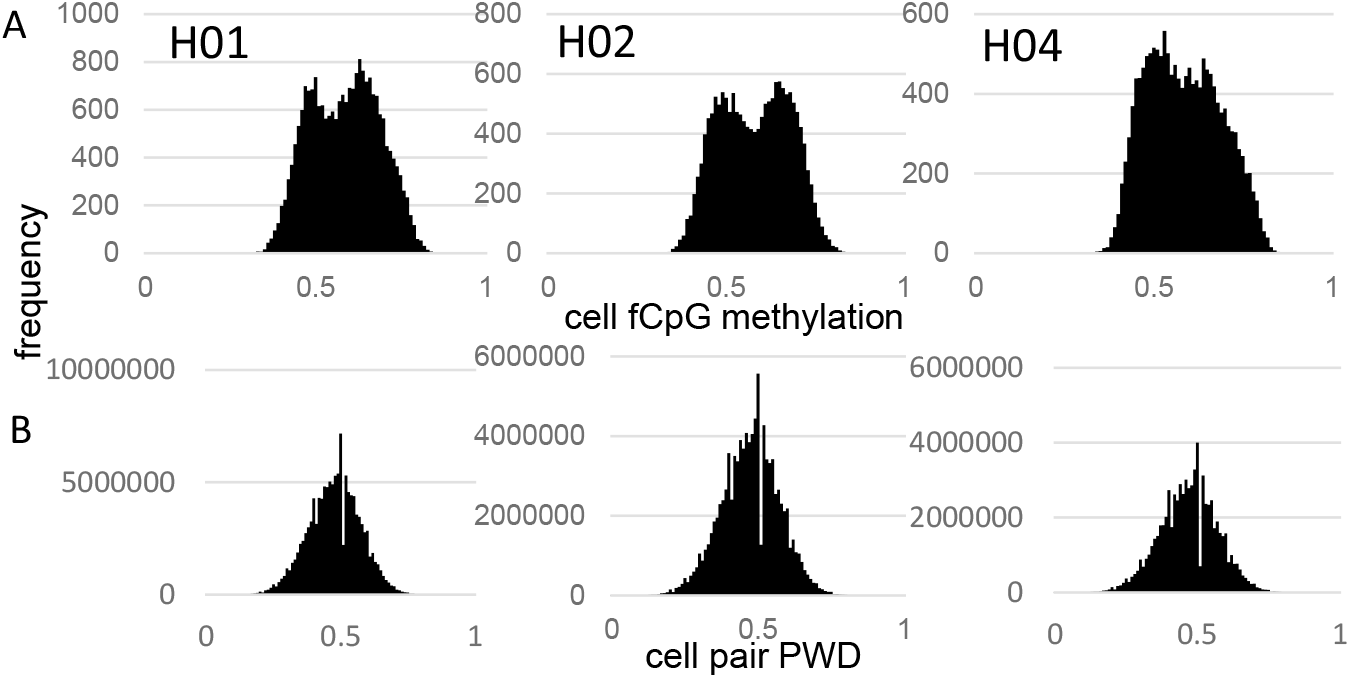
fCpG barcode methylation. **A)** Barcode methylation was variable between cells with averages ∼ 50% **B)** Most cells had different barcodes with average PWDs ∼ 0.5.

### fCpG barcodes initialize methylated and change with cell division

fCpG barcode patterns were similar between the brains, and data are presented for H01, with H02 and H04 presented in Figs S1 and S2. Methylation was variable between cells of the same type and average methylation was highest in the pons (PN) and thalamus (THM, intermediate for other inhibitory neurons, and lowest for excitatory neurons, glial cells, and cerebellar cells (Fig 2A). Outer layer cortical excitatory neurons (L2_3) that are made later during development were less methylated than inner cortical excitatory neurons (L4_6) that appear earlier. Predominately methylated individual fCpG sites were common in hindbrain neurons (PN, THM), less frequent in other inhibitory neurons, and rare in excitatory neurons (Fig 2B).

**Fig 2:**
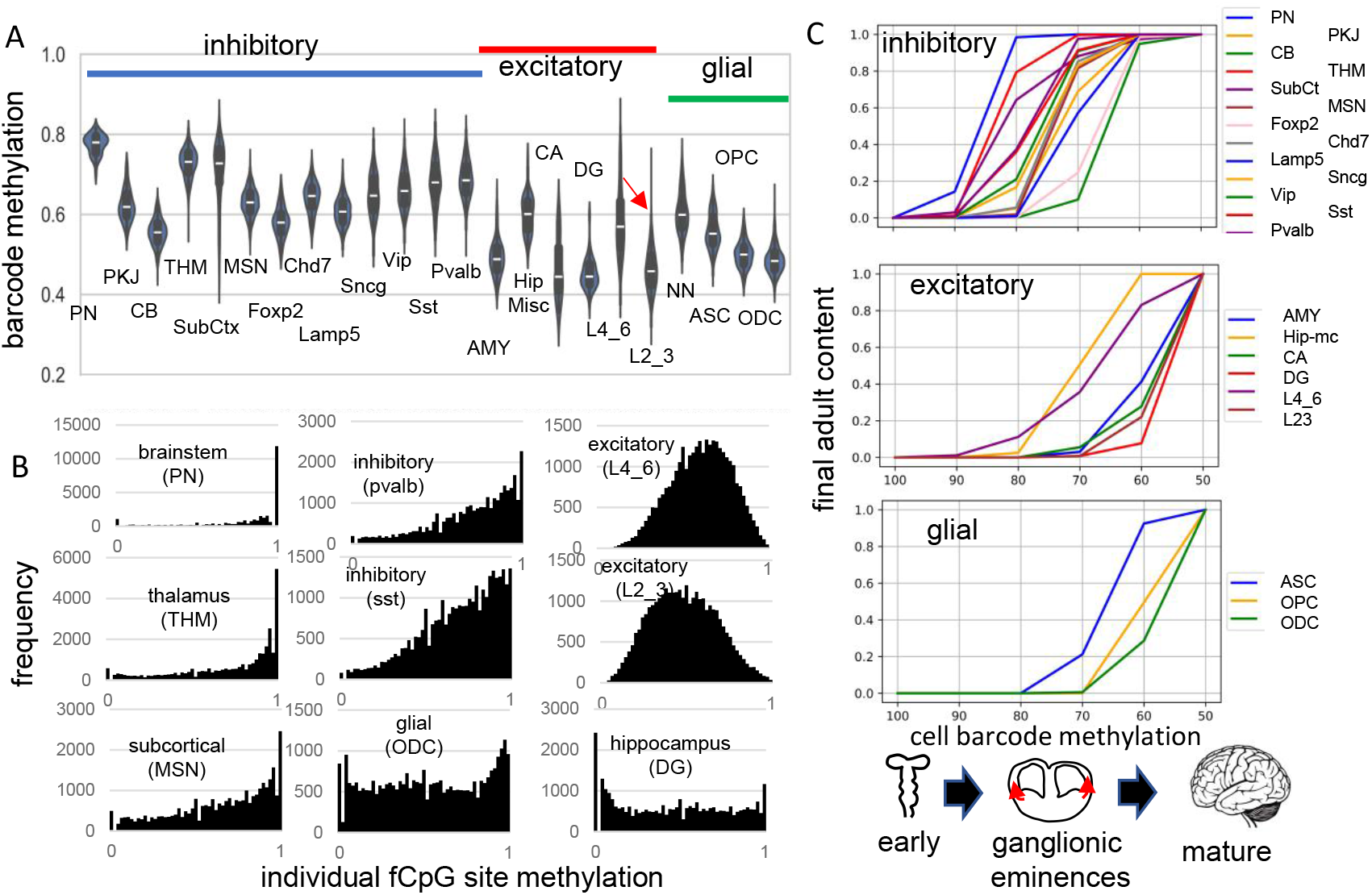
Higher fCpG barcode methylation in earlier emerging cells. **A)** Average barcode methylation was higher in the brainstem and inhibitory neurons. Barcode methylation was lower for excitatory neurons, cerebellar, and glial cells. Notably, average methylation was lower for outer cortical (L2_3) compared to earlier appearing inner (L4_6) cortical excitatory neurons. Abbreviations are as in reference 3, with L2_3 all outer and L4_6 all inner cortical excitatory neurons, and NN are non-neuronal cells other than ASC, OPC and ODC. **B)** Most fCpGs appear to start methylated in a progenitor because nearly all individual fCpGs are methylated in inhibitory neurons in the subcortex (PN, THM, MSN). Many fCpGs in inhibitory neurons (pvalb, sst) are still predominately methylated. Few fCpGs in excitatory neurons that differentiate later in development are fully methylated. Glial cells that also emerge late in development, and hippocampal cells that may divide postnatally had variable methylation with both highly methylated and unmethylated fCpGs. **C)** Barcodes are assumed to become fixed when their cells stop dividing and differentiate. Therefore, barcode methylation levels can indicate when neurons emerge during development, and can be correlated with a cartoon of physical caudal to rostral brain development. The X axis indicates the barcode methylation of individual cells and is assumed to roughly correlates with calendar time. The Y axis indicates the cumulative proportion of cells of each type present at each methylation level. A value of 0 indicates that cells of given type are not yet present and a value of 1 indicates the adult content of this cell type has been reached. At the start of development, inhibitory neurons (PN) in the pons with highly methylated barcodes appear first. More inhibitory neurons, made in the ganglionic eminences, appear and reach their final adult contents before many cortical excitatory neurons and glial cells appear. Notably, barcode methylation indicates many lower cortical layer neurons appear earlier in life relative to outer cortical neurons that reach adult levels late in development. Brain contents inferred by adult barcodes may differ from actual neonatal brains because neurons that die during development are not sampled in adult brains.

The methylation hierarchy is consistent with a barcode initialized with predominately methylated fCpGs in a brain progenitor cell. Barcodes becomes progressively demethylated and are fixed when their cells stop dividing and differentiate, which occurs at different times and places during brain development. Barcodes for each cell type had a range of methylation (Fig 2A), consistent with synchronous development rather than a strict stepwise process. Very simplistically, the hindbrain with mature neurons (6) forms early in development, followed by inhibitory neurons in the ganglionic eminences, and then excitatory neurons and glial cells in the cortex. Barcode methylation follows this temporal development and reconstruct when specific neuron types start to appear and reach their adult contents (Fig 2C). For example, after barcodes change from ∼100 to ∼70% methylated, most adult brainstem (PN) and inhibitory neurons are present but excitatory neurons are fewer, with very few adult outer layer (L2_3) neurons. Outer excitatory and glial progenitor cells are present (7), but their barcodes continue to demethylate until they stop dividing and differentiate later in development. This stopwatch like pattern, with more demethylated barcodes in later appearing cell types, was present in all three adult brains.

### fCpG barcodes are polymorphic

A progenitor cell barcode is assumed to become increasingly polymorphic with subsequent divisions. This pattern was observed, with average barcode PWDs lowest in brainstem cells (PN), intermediate between other inhibitory neurons, and highest for excitatory neurons (Fig 3A). Most cells had different barcodes, with an overall average PWD of ∼0.47 (Fig 1B). Cells of the same phenotype were more similar with lower average PWDs (Fig 3A, 3B), suggesting they are more related to each other and have common progenitors.

**Figure 3:**
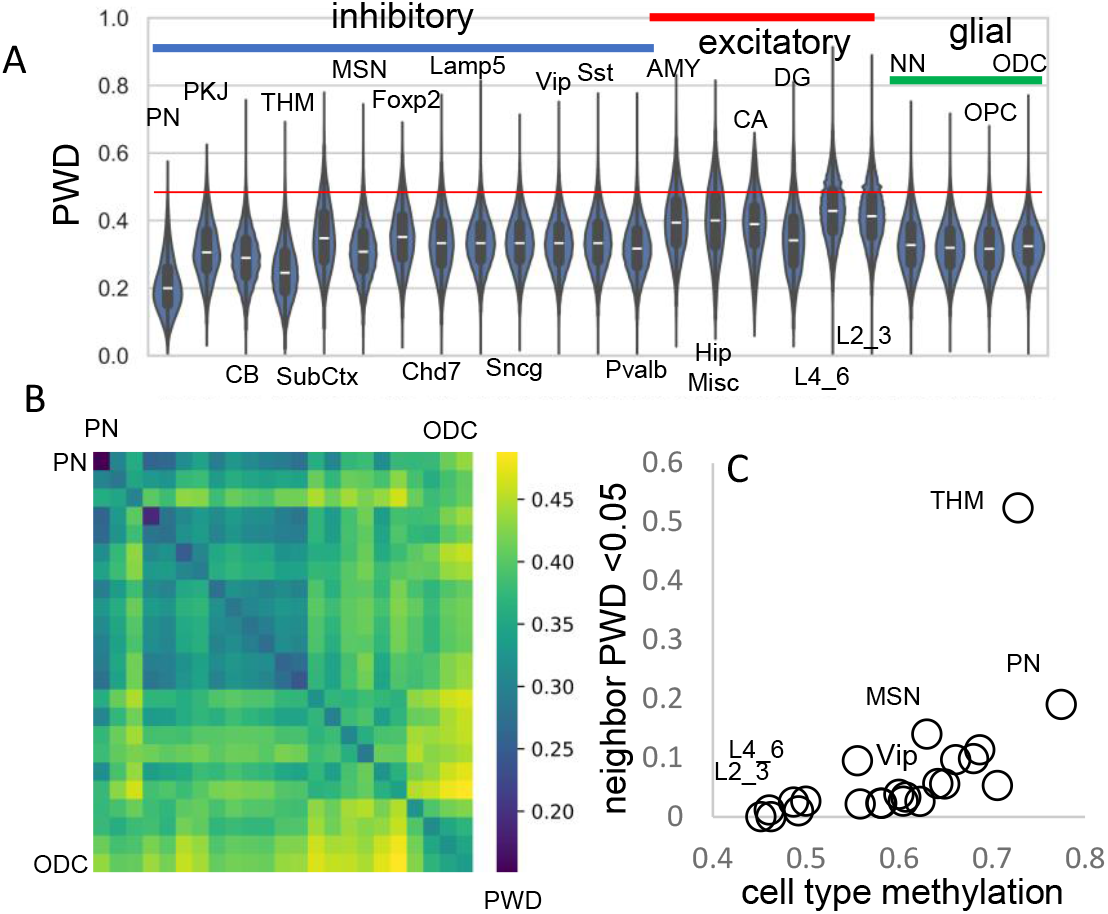
Related cell pairs. **A)** Most cells had different barcodes with average PWDs between cell pairs of ∼ 0.5. Cell pairs of the same phenotype had different barcodes but were on average more related to each other. **B)** Heatmap showing that cells of the same phenotype are more related. **C)** Cells that emerge early in development are more related and more methylated. Closely related nearest neighbors (PWD <0.05) are numerically more common for more methylated cell types.

The human brain has billions of cells and relatively few cells were sampled from each region. Consistent with sparse sampling, cell pairs with nearly identical barcodes and smaller PWDs (< 0.05) were rare. To help distinguish between ancestry and chance, cells within and between brains were compared (Table 1). Closely related cell pairs were ∼2.9 times more frequent within a brain (average ∼9.8 per million) compared to between brains (average ∼3.4 per million). Closely related cells had fewer matching fCpG sites (∼35 compared to ∼48 for all cell pairs) and were more common early in development when barcodes are more methylated (Fig 3C), indicating that lower barcode complexity favors matching. Overall, fCpG barcodes are sufficiently polymorphic to distinguish between most adult brain cells.

### Brain lineage trees

It should be possible to reconstruct human brain development if barcodes record ancestry. fCpG barcodes from ∼1,000 brain cells with different phenotypes yield trees with a standard phylogeny software that resemble caudal to rostral development (Fig 4). The trees are rooted by a progenitor with a fully methylated barcode, and branches progressively yield brainstem neurons (PN), a subset of excitatory lower (L4_6) neurons, thalamic neurons (THM), inhibitory neurons, cerebellar cells, and glial cells. Excitatory neurons branch last, and hippocampal neurons (CA, DG) that may divide postnatally (8) were at the terminus. Cells generally grouped by phenotype, with some early appearing excitatory neurons admixed among inhibitory neurons. Similar trees were observed for H02 and H04, albeit with less separation between inhibitory and excitatory neurons for H04 (Fig 4A). Barcode lineage trees are largely consistent with expected sequential neuronal differentiation.

**Figure 4:**
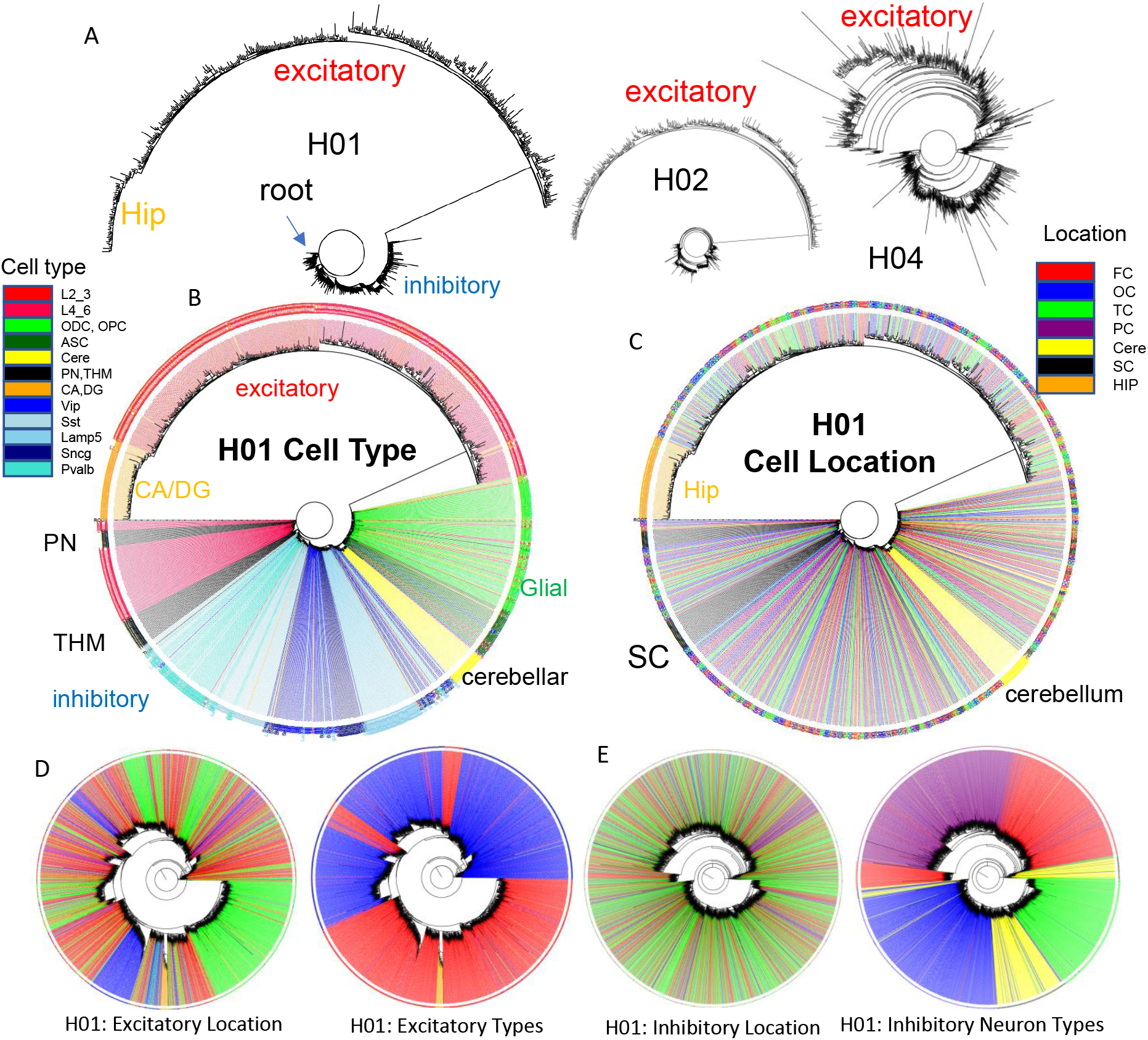
Brain trees. **A)** Barcodes from 960 cells form trees using IQtree (19) that are rooted by a fully methylated progenitor, and generally follow caudal to rostral brain development, with sequential branching of inhibitory neurons, cerebellar neurons, and excitatory neurons, with hippocampal neurons furthest from the start. Trees are similar between the brains, with H04 inferring less distance between inhibitory and excitatory lineages. The trees illustrate the ability to produce phylogenies with IQtree, but the phylogenies are limited by sparse cell sampling and that barcodes may be similar by chance. The degree of confidence was generally low, with bootstrap branch support typically less than 15%. **B)** H01 tree with labeled cell types. Neuron types generally clustered by phenotypes with closely branching excitatory and inhibitory neurons more common earlier in development. **C)** H01 tree with labeled cell locations. Related inhibitory and excitatory neurons can be found in different parts of the brain (FC = frontal (red), TC = temporal, OC = occipital (blue), PC = parietal, HIP = hippocampus (orange), cere = cerebellum (yellow), SC = subcortical (black). **D)** H01 tree with ∼2,853 cortical excitatory neurons has more evidence of localized radial migration because related neurons are more often found in the same cortical region. Excitatory neurons cluster by subtype, and closely related lower and upper excitatory neurons were still few. **E)** H01 tree with ∼2,847 cortical inhibitory neurons still retains evidence of tangential migration with related neurons scattered throughout the cortex. Inhibitory neurons cluster by subtype with switching between some closely related pairs.

### Cell lineage fidelity and cortical migration

Uncertain for mouse and human development is whether inhibitory and excitatory neurons originate from shared or distinct progenitors (3,9,10). Barcodes could potentially record neuronal differentiation patterns and lineage fidelity can be quantified by comparing most closely related cells or nearest neighbor cell pairs with PWDs < 0.05. The approach remains speculative due to several factors: the absence of direct experimental validation, limited experimental cell sampling, and the possibility that barcode similarities may arise by chance rather than reflecting true biological relationships. Lineage fidelity was high (>90%) for inhibitory neurons (Fig 5A). Excitatory lineage fidelity was slightly lower, indicating that some excitatory and inhibitory neurons may share common progenitors (10). Lineage trees indicate common progenitors are present earlier in development, and excitatory neurons that appear later do not have many closely related inhibitory neighbors (Fig 4B). The barcodes documented the known switching between inhibitory neuron subtypes (Fig 5B). Hindbrain, excitatory, and non-neuronal cells had more subtype lineage fidelity.

**Figure 5:**
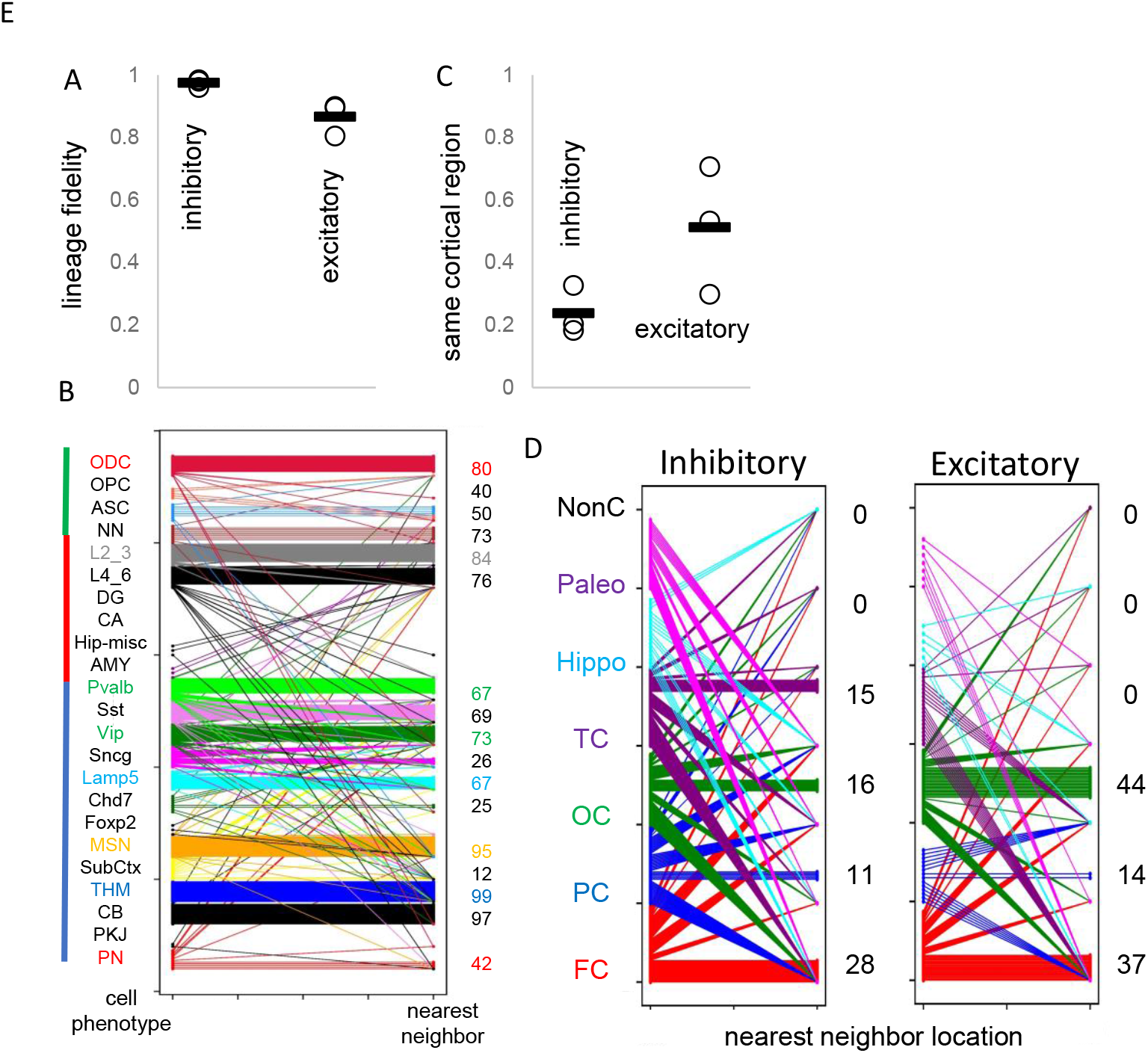
Lineage fidelity, migration, and differentiation. **A)** Inhibitory neurons have higher lineage fidelity because nearest neighbor pairs (PWD <0.05) were nearly always both inhibitory neurons. Excitatory neurons had slightly less lineage fidelity because a nearest neighbor was more often an inhibitory neuron. Data are for all three brains. **B)** Nearest neighbor inhibitory neuron pairs often had subtypes differences. More lineage fidelity was generally present for subcortical and excitatory neurons. Numbers indicate percent lineage subtype fidelity. **C)** Nearest neighbor inhibitory and excitatory neuron pairs showed evidence of tangential migration because they were found in different cortical regions. The data indicate greater evidence of inhibitory neuron tangential migration than for excitatory neurons. However, the extent of migration is uncertain because of sparse sampling and because barcodes may be similar by chance. Data are for all three brains. **D)** Nearest neighbor neurons were scattered in the cortex. Numbers indicate percent location fidelity. (NonC = non-cortical location, Paleo = paleocortex)

Barcodes can also potentially infer migration because their neurons are annotated by their adult locations. Daughter cells with similar barcodes could be sampled from the same region, or from different regions if migration occurred. Trees (Fig 4C) indicate that most neurons sampled from the brainstem and hippocampal regions are related and localized to their respective regions. Inhibitory neurons were scattered throughout the cortex, consistent with their differentiation in the ganglionic eminences and subsequent tangential migration to the cortex.

Nearest neighbor inhibitory cortical neuron pairs were found in the same cortical region ∼25% of the time (Fig 5C and 5D). Nearest neighbor excitatory neuron pairs were also scattered throughout the cortex, but less than inhibitory neurons, and were in the same cortical region ∼50% of the time. The barcode data indicate more inhibitory rather than excitatory neuron tangential migration, but the extent of migration is uncertain due to sparse sampling and because barcodes can match by chance.

The poor ability to detect localized excitatory neuron radial cortical migration with ∼1,000 cell whole brain trees (Fig 4C) may reflect that sparse sampling is unlikely to include multiple neurons from the same small clonal region that originated from common subventricular progenitors (radial unit hypothesis (11)). Greater localized excitatory neuron migration was seen when trees were reconstructed with more (∼2,800) neurons, while inhibitory neurons still showed scattered tangential migration (Fig 4D). Neurons of the same subtype were still more related. Hence, lineage trees appear to increase their resolution with more cells, albeit related lower and upper excitatory neuron pairs were still uncommon, which may reflect the unlikely chance of sampling very small radial clonal units.

## Discussion

fCpG barcodes are potential markers of somatic cell ancestry and not cell type classifiers, although cells of the same phenotype are often related because they originate from common progenitors (see new Supplement for fuller discussion). Dynamic barcodes would be useful to study human tissues, but testing their performance is difficult. Ideally, samples obtained at different times would document how they change. The brain facilitates barcode validation because it periodically stores neurons that stop recording at relatively defined times and locations (1,13). Specific neuron subsets recovered from the adult brain allow for sampling through time and before birth.

This serial sampling strategy facilitated fCpG barcode validation. The barcode appeared to start predominately methylated in multiple individuals and became sufficiently polymorphic to distinguish between thousands of neurons. Barcode changes appear to represent replication errors because they reconstruct lineage trees roughly consistent with caudal to rostral brain development. Barcode methylation may indicate when different neurons that survive to adulthood appear in the neonatal brain (Fig 2C).

The current barcode indicates that most inhibitory and excitatory neurons have relative distinct progenitors, consistent with the lineage dendrograms reconstructed with neuron specific methylation of the same data (3). There was also evidence for common inhibitory and excitatory progenitors (10), primarily for earlier emerging excitatory neurons. Tangential migration was also detected, manifested by inhibitory neurons with closely related barcodes in different cortical regions. Tangential excitatory neuron migration was also detected, albeit related excitatory neurons were more localized than inhibitory neurons. Tangential migration is also seen with sequencing studies that find neurons with specific mutations in multiple brain regions (14-17).

fCpGs more efficiently distinguish between cells than mutations due to higher replication error rates. Although average methylation decreases with time, both demethylation and remethylation are likely because fully demethylated neurons were not observed, and balanced fluctuating methylation is inferred in other tissues when CpG sites are ∼50% methylated in bulk tissues (2). More adult divisions in brain cancers did not saturate the barcode with average fCpG methylation ∼50% (Fig S3). Fluctuating methylation complicates lineage tracing but backmutations can be modeled for ancestral reconstructions. Lineage resolution could be improved by combining mutations and fCpGs.

Weaknesses of this study including very sparse cell sampling and lack of uniform CpG sites comparisons between neurons. Like many human fate markers studies, it is difficult to independently verify accuracy. Interpretation of the barcodes relies on several untested assumptions including relatively constant error rates between fCpG sites and through aging, neutrality, and a predominately fully methylated start in the progenitor cell. Epigenetic remodeling occurs after progenitors stop dividing (12), which could erase ancestral barcode information. However, neurons of the same type were both closely and remotely related, indicating that such epigenetic remodeling does not systematically alter the fCpG sites. In addition, fCpG barcodes appear to be relatively stable through aging (new Supplement). Inferred lineage trees (Fig 4) had relatively low statistical support for their branches and are presented to demonstrate that the barcodes are readily organized into trees with a commonly used phylogeny software.

Technical improvements such as targeted bisulfite sequencing of a limited number of informative fCpGs could lead to more consistent coverage and less expensive sequencing of more neurons. Single cell measurements of small numbers of fCpGs, and snMCode cell type specific CpG sites (3), could efficiently reconstruct human brain lineages, although back changes add complexity. A barcode of 100 fCpGs has enough complexity (2^100^ or ∼1×10^30^) to potentially distinguish between most excitatory neurons, with less resolution early in development when cells are inherently more related.

The analysis of more brains can verify that a fCpG barcode starts predominately methylated in most individuals. A common initialized state could facilitate standardized human fate maps and comparisons between individuals. Many polymorphisms linked to brain abnormalities such as autism are in neuronal proliferation, migration, and maturation pathways (18), and this preliminary survey indicates lineage heterogeneity between individuals (Fig 4A). fCpG barcodes have been applied to the intestines, endometrium, and blood (2), and could be found for multiple other tissue types, helping to unravel human development and aging.

## Methods

### Brain Single Cells

Single cells with their annotations and methylation at each fCpG site were read from single cell files downloaded from GEO (GSE215353) and supplemental files from reference 3. Lists of fCpG sites and data summarized for the Figures are in Supplemental File 2. The cells and methylation at the fCpG sites are in Supplemental Files 3-5. PWDs were calculated between all cell pairs with at least 30 matching fCpG sites, with PWD data matrices in Supplemental File 6. Additional details are provided in the new Supplement.

### IQtree

IQtree (19) tree was downloaded and run on a server with 64 cpus and 32 GB of memory. The model (GTR2+FO+G4) accounts for backmutation and binary data with missing values. Bootstraps were 1,000 per tree with 3,000 iterations for whole brain (∼1,000 cells) trees and 1,000 iterations for inhibitory or excitatory (∼2,800 neurons) trees. Trees (.treefile) were displayed with FigTree (http://tree.bio.ed.ac.uk/software/figtree/) with truncation of long branches (generally fewer than 10) for display purposes. The cells used for the trees are in Supplemental File 7.

## Supporting information

New Supplement

## Acknowledgements

This work was supported by grants from the NIH (P01CA196569 and CA271237). I thank Drs. Trevor Graham and Heather Grant for useful discussions, and Omar Khan and Nikhil Krishnan for initial studies. The author thanks all of the researchers that helped produced the high quality very valuable and freely available data used for analysis.

---

**Supplementary Figure 1:**
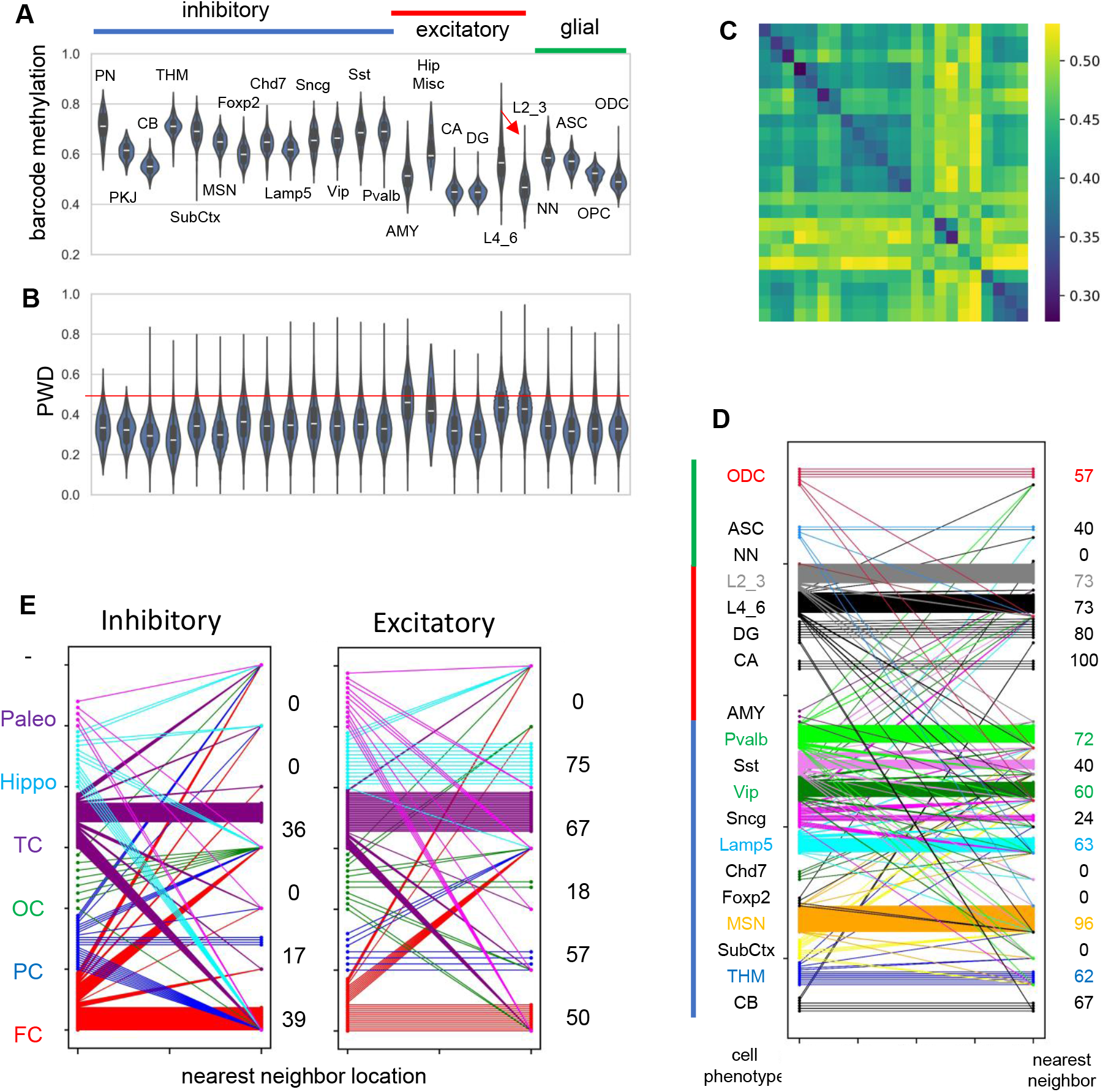
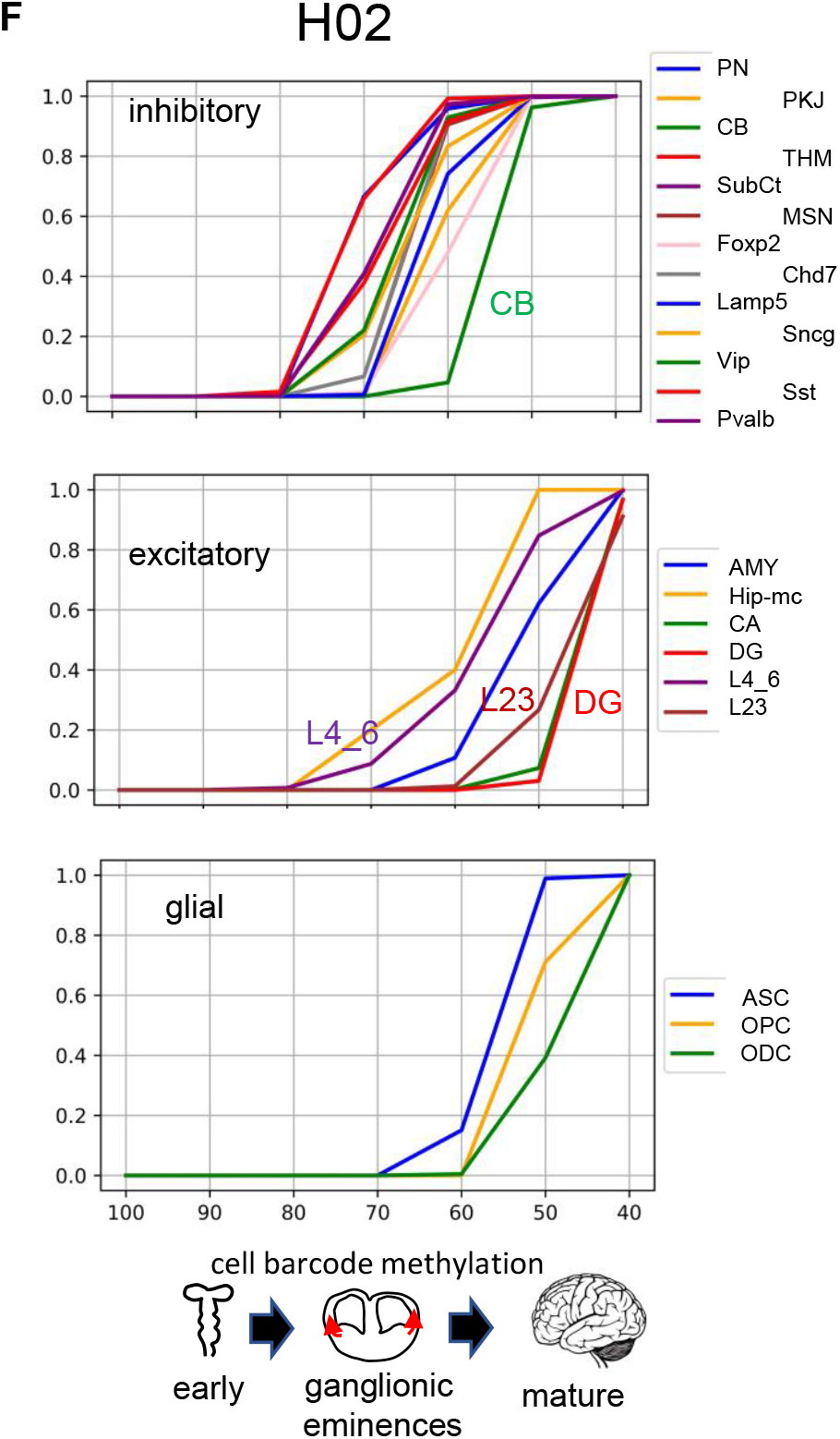
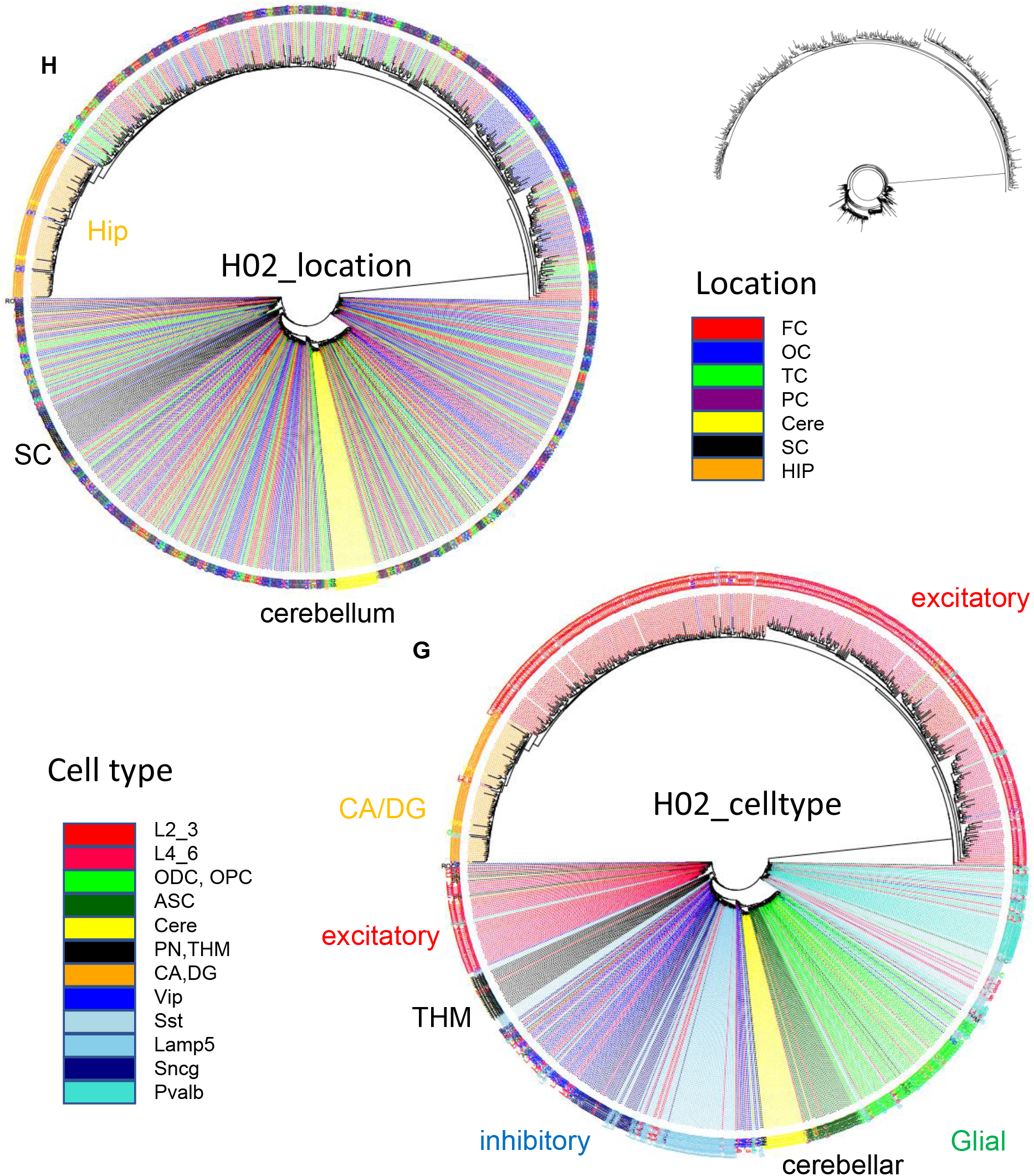
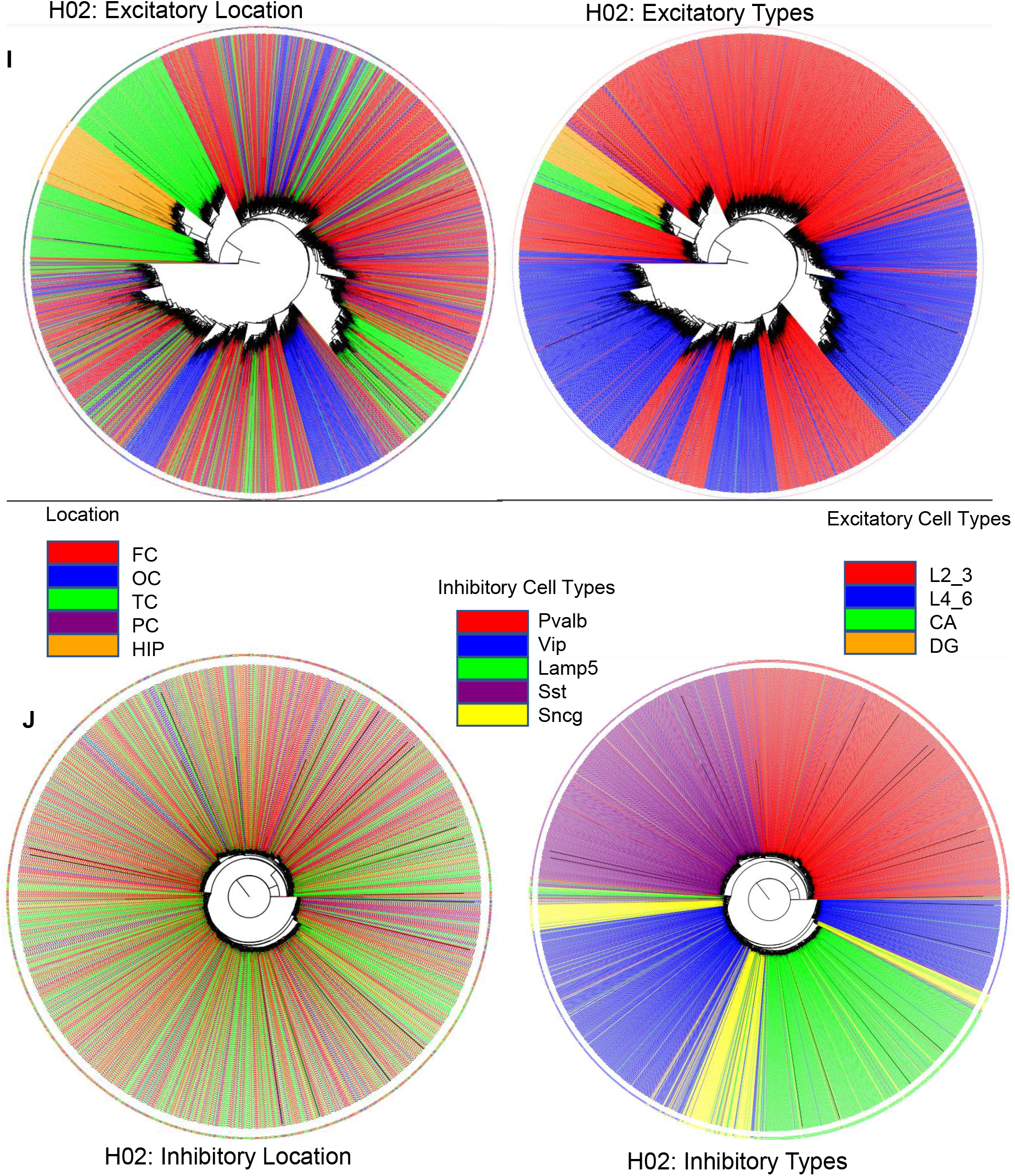
H02 data. **A)** Barcode methylation for different cell types **B)** PWDs between cells of the same type **C)** PWDs between cell types **D)** Lineage cell type fidelity between nearest neighbor pairs (PWD<0.05) **E)** Location fidelity between nearest neighbor pairs (PWD<0.05) **F)** Barcode methylation versus final adult brain content indicates that inhibitory neurons appear first and reach their adult levels before excitatory or glial cells. **G)** Ancestral tree with 1,001 cells rooted at a fully methylated progenitor shows sequential branching with excitatory, then brain stem, inhibitory and cerebellar neurons, then glial cells, and finally excitatory neurons with hippocampal neurons at the end. **H)** Related cells colocalize for brain stem, cerebellar, and hippocampal neurons. Inhibitory neurons are more scattered. Excitatory neurons are also scattered with some localization within the cortex (see I & J for trees with more neurons) **I)** Ancestral tree with more (2,806) excitatory neurons shows more localization between related neurons in the hippocampus, occipital and temporal cortex. Related neurons also tend to have similar phenotypes. **J)** Ancestral tree with more (2,788) inhibitory neurons still shows scattering between related neurons. Related neurons tend to have similar phenotypes.

**Supplementary Figure 2:**
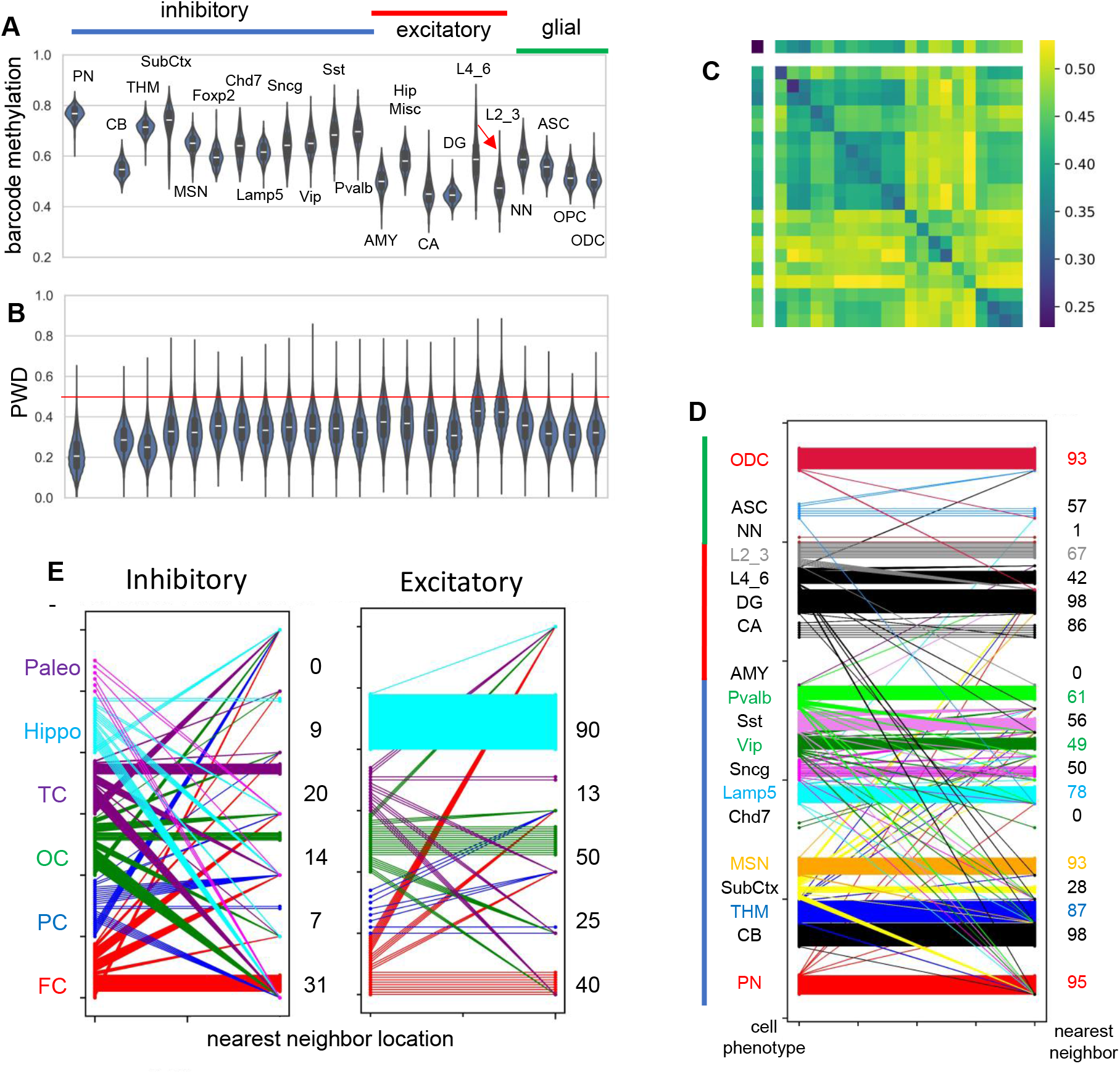
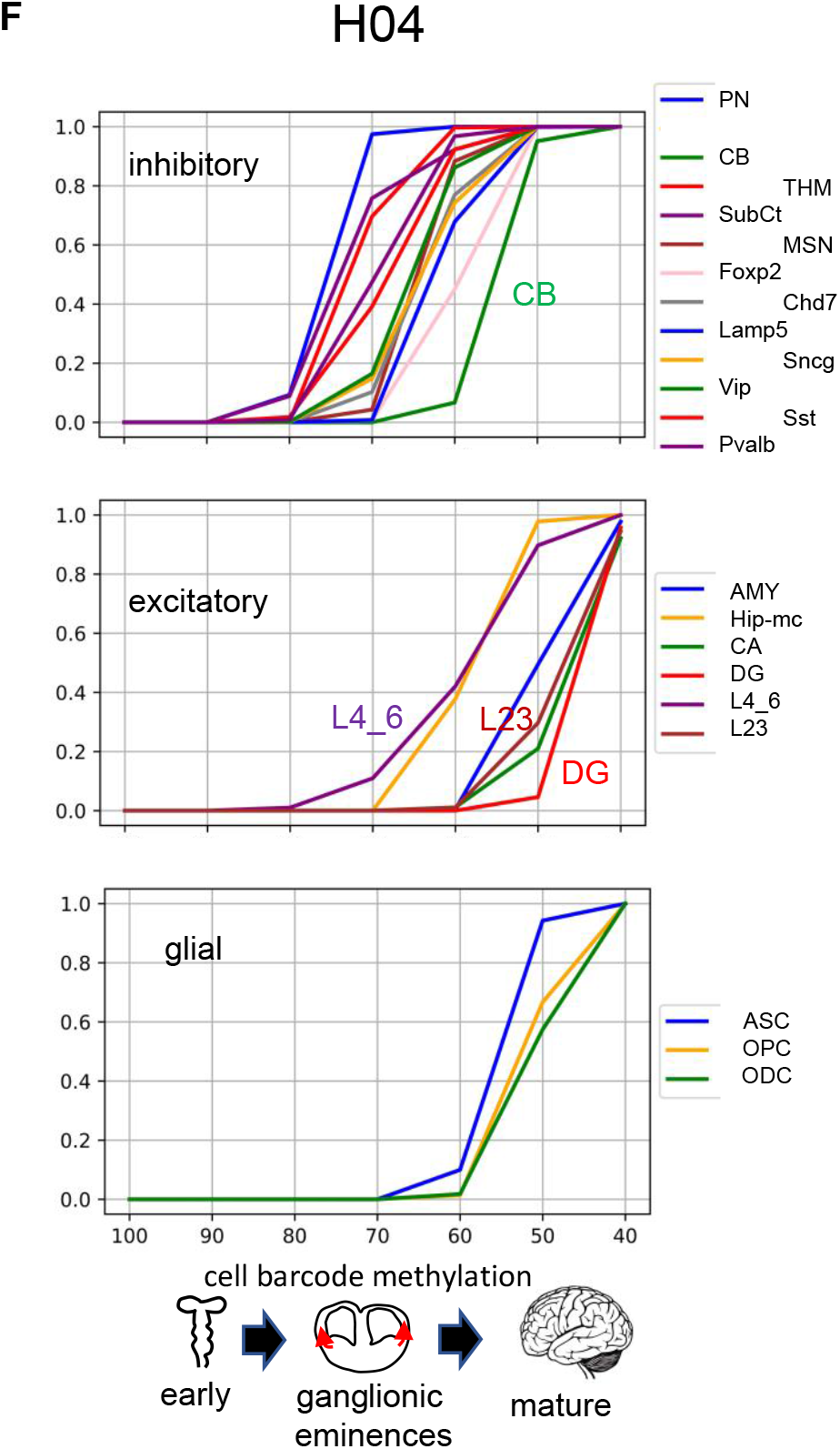
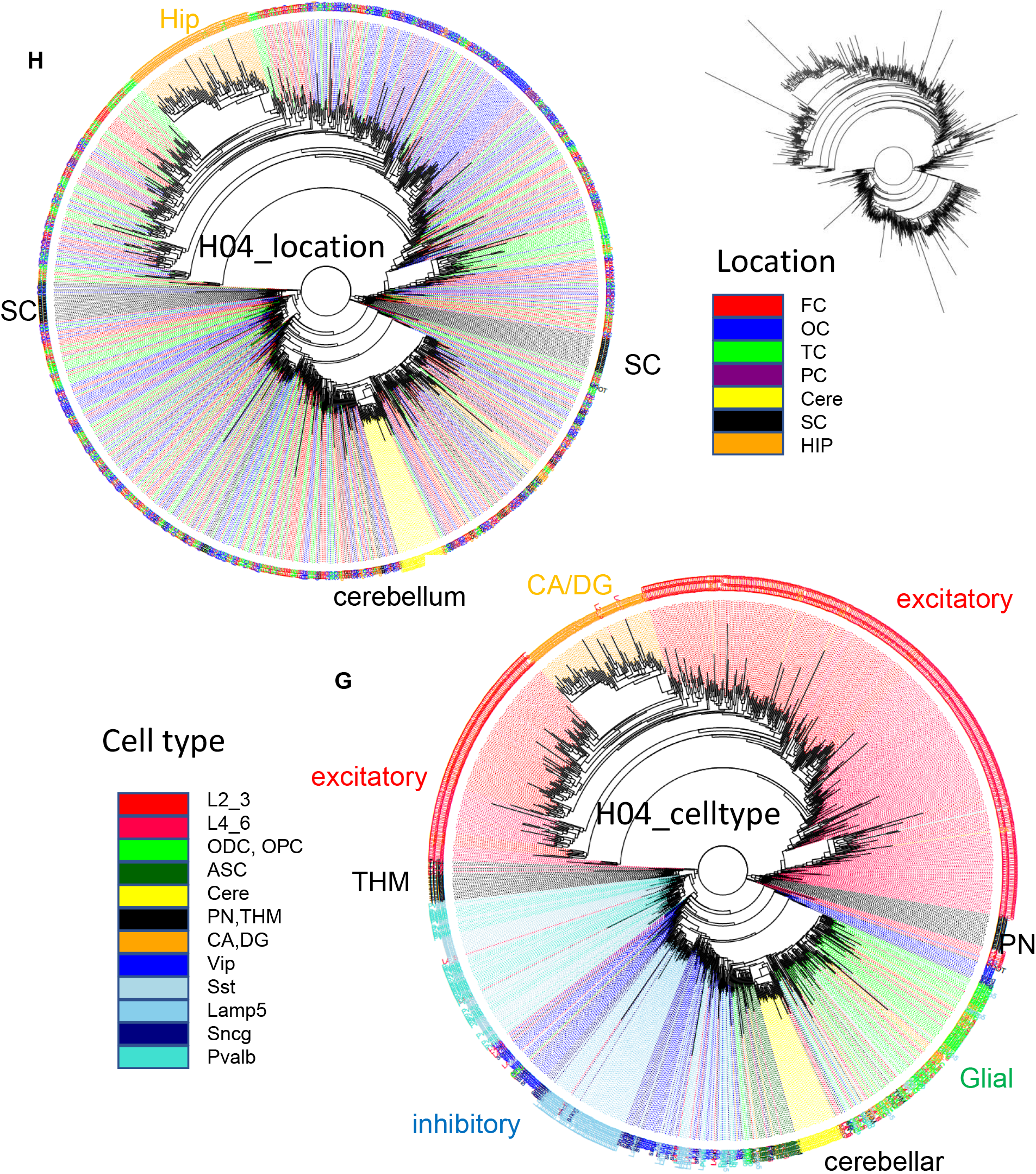
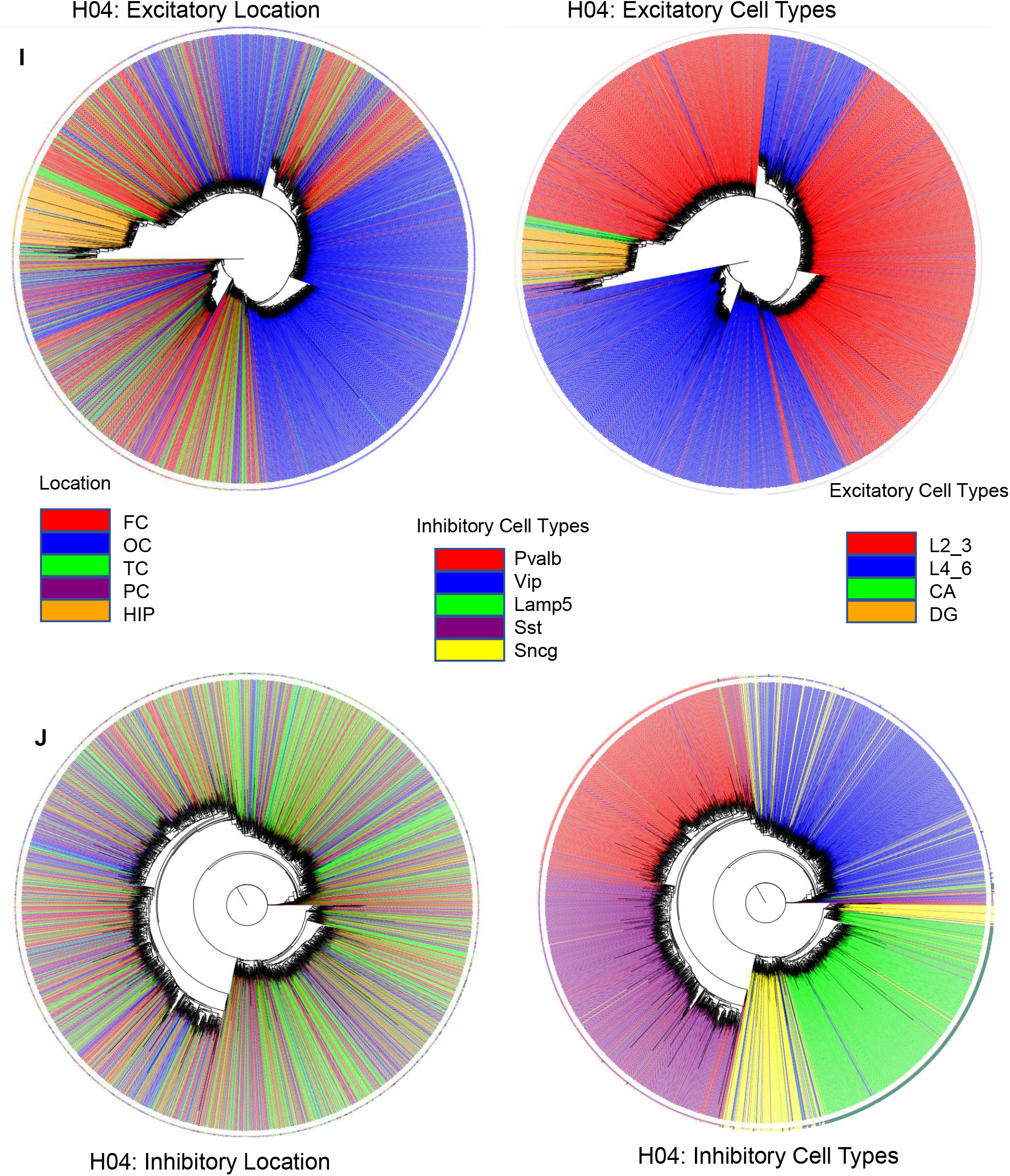
H04 data. **A)** Barcode methylation for different cell types **B)** PWDs between cells of the same type **C)** PWDs between cell types **D)** Lineage cell type fidelity between nearest neighbor pairs (PWD<0.05) **E)** Location fidelity between nearest neighbor pairs (PWD<0.05) **F)** Barcode methylation versus final adult brain content indicates that inhibitory neurons appear first and reach their adult levels before excitatory or glial cells. **G)** Ancestral tree with 1,033 cells rooted at a fully methylated progenitor shows sequential branching with excitatory, then brain stem, inhibitory and cerebellar neurons, then glial cells, and finally excitatory neurons with hippocampal neurons at the end. **H)** Related cells colocalize for brain stem, cerebellar, and hippocampal neurons. Inhibitory neurons are more scattered. Excitatory neurons are also scattered with some localization within the cortex (see I & J for trees with more neurons) **I)** Ancestral tree with more (2,797) excitatory neurons shows more localization between related neurons in the hippocampus, occipital and temporal cortex. Related neurons also tend to have similar phenotypes. **J)** Ancestral tree with more (2,752) inhibitory neurons still shows scattering between related neurons. Related neurons tend to have similar phenotypes.

