## Supplementary material for "Human Brain Ancestral Barcodes": New Supplement

The purpose of this supplement is to address the broad concerns raised by the three reviewers. The goals are to better explain the mechanics of fCpG barcodes, present new data illustrating their reproducibility and stability with aging, and demonstrate how barcodes can record ancestry after conception, even in the context of active demethylation.

**Differences Between fCpGs and Traditional Cell Type Classifiers**

Both fCpG barcodes and traditional DNA methylation-based cell type classifiers can differentiate between cells, but their identification processes and underlying biological principles differ significantly. Traditional cell type classifiers focus on identifying CpG sites that are consistently differentially methylated across distinct cell types. Success in this approach yields a barcode, such as the snMCode CpG sites (1), where each cell of a given phenotype shares the same methylation pattern, distinct from those of other phenotypes.

In contrast, the identification of fCpGs is fundamentally different because fCpG methylation can vary among cells, even within the same type. In traditional screening for cell type classifiers, a CpG site is retained if its methylation is consistent across cells with the same phenotype and discarded if it varies. Conversely, when screening for fCpG sites, a CpG site is retained if its methylation differs among cells and discarded if it skews toward either methylated or unmethylated states. The fCpG barcode captures random errors accumulated throughout development, recording ancestry, while the methylation profiles used in cell type classifiers more accurately reflect terminal differentiation.

**Lineage Tracing: Cladistics and Ancestry**

Although fCpG selection and cell type classifier selection are fundamentally opposed, their barcodes may align if clades of cells sharing the same phenotypes arise from common progenitors (Fig 1A). This study supports the notion that neurons of the same type typically originate from common progenitors, as cells within a clade generally exhibit polymorphic yet more similar fCpG barcodes than those from different clades. fCpG barcodes may provide complementary insights into cell mitotic ages, diversity within a clade, and migration patterns of daughter cells—factors that are often more challenging to discern from RNA-seq data alone.

**fCpG Barcode Mechanics**

The fCpG barcode begins as predominately methylated, reflecting the observation that inhibitory neurons in the pons—emerging early in development—are predominantly methylated across the three brains examined. It is assumed that random errors occur when the barcode is replicated during cell divisions, as illustrated in a cartoon (Fig 1B). As errors are perpetuated and new ones accumulate, closely related cells tend to exhibit more similar methylation patterns than those that are less related. The differences among these patterns can be quantified by calculating the average pairwise differences (PWDs) between barcode fCpGs.

The barcode mechanics are more formally described by simulations that match the experimental data. These simulations start with a single cell possessing a fully methylated barcode. Key parameters include fCpG methylation error rates, the numbers of cell divisions, and whether a cell division yields two, one, or zero daughter cells. A simulation that broadly matches the experimental data for excitatory neurons has 150 cell divisions and a fCpG error rate of 0.01 per division, applied equally to both methylated-to-unmethylated and unmethylated-to methylation flips (Fig 2).

Notably, even with a high error rate of 0.01 per division ancestry may be reconstructed because each fCpG sites experiences relatively few changes or methylation flips. By the end of 150 divisions, the average methylation level approaches approximately 50%, with most fCpG sites experiencing three or fewer changes in methylation. Among the methylated sites, around 20% remain unchanged (methylated), while approximately 25% of fCpG sites have undergone two flips (from 1 to 0 to 1). For unmethylated sites, about one-third experience one flip (from 1 to 0), and roughly 15% have had three flips (from 1 to 0 to 1 to 0). Only about 5% of sites experience more than three flips. Despite this, the randomness of these flips leads to highly polymorphic barcodes among the final cells, with more related cells having more similar barcodes. As noted by Reviewer 1, this barcode mechanism is essentially a mitotic clock that becomes polymorphic.

**Simulations of Neurogenesis**

Simulations that align with experimental data can effectively illustrate the mechanics of the barcode. The first simulation, designed to model early neurogenesis in the hindbrain, features a limited number of divisions and a basic exponential expansion, beginning with a single cell possessing a fully methylated barcode (Fig 3A). After 19 divisions, resulting in ~500,000 cells, the average methylation across the population declines to approximately 85%, while the average PWD between cells increases to about 0.28. The proportion of closely related neighboring cells (with PWDs < 0.1) progressively decreases to around 45%.

The simulation of inhibitory neurogenesis extends to 50 divisions, also starting from a single cell with a fully methylated barcode (Fig 3B). In this scenario, cell division is balanced by cell death to maintain a small population early in development, followed by subsequent expansion. By the end of 50 divisions, average cell population methylation falls to about 70%, and average PWD increases to around 0.35, with the proportion of closely related neighboring cells dropping to approximately 20%.

The simulation of excitatory neurogenesis involves 150 divisions, where early development similarly balances cell division and cell death, followed by expansion (Fig 3C). The average cell population methylation decreases to about 55%, and the average PWD among cells rises to around 0.4, with the proportion of closely related neighboring cells declining to about 5%. While these simulations do not fully capture the complexity of neurogenesis such as the variation in differentiation times, they illustrate how fundamental barcode mechanics can yield neuron populations broadly consistent with experimental data.

**Barcode Cell Division Dynamics During Early Development**

The public review requested additional experimental data, and Reviewer 1 raised concerns that active methylation processes early in development could obscure any signature of inaccurate fCpG methylation. Direct comparisons of immediate human daughter neurons pose challenges, and it is uncertain what a relevant cell culture study might be.

Whole genome bisulfite single-cell data are available for human germ cells, zygotes, 2 to 8-cell embryos, morulae, and ICM samples (2). The data show that barcodes are more methylated in germ cells and gradually become less methylated as development progresses to the ICM (Fig 4). While fCpG barcodes are polymorphic between unrelated samples, they exhibit greater similarity among closely related cells in the 2 to 8-cell stage, with reduced similarity in morulae and ICM samples (Fig 4).

This additional data indicates that more closely related daughter cells have more similar barcodes, even amidst the chaotic background of early development, where both active and passive demethylation largely erases germ cell methylation (3). Notably, although a neurogenic lineage is present among the embryonic cells, none of the cells displayed fully methylated barcodes, suggesting that the anticipated remethylation in a neurogenic progenitor has not yet occurred. The activation of DNMT3A during early neurogenesis (4) may ultimately lead to the establishment of fully methylated barcodes that subsequent become polymorphic.

**Neuron Barcode Reproducibility And Stability With Aging**

New single-cell neuron datasets (5,6) became available while the manuscript was under review. The core concept behind brain ancestry barcodes is that they can be used to compare different brains, assuming all brains start with a fully methylated barcode and share similar developmental trajectories. These new datasets offer valuable opportunities to test for barcode reproducibility and stability. One dataset (Fig 5) measured neurons in a seven-month-old brain from the frontal cortex and hippocampus (5). The adult fCpG barcode methylation levels observed are largely present in the infant brain, supporting the idea that fCpG methylation patterns are established prenatally through replication errors and remain stable once cell division ceases.

A second dataset (6) sampled the same region of the frontal cortex (Brodmann area 46) from young (~25 years) and old (~70 years) brains. This study did not reveal extensive changes in DNA methylation with aging but identified a small number of differentially methylated CpG regions (6). Consistent with the overall stability of DNA methylation during aging, average barcode methylation (Fig 6A) and PWDs (Fig 6B) among the ten samples from six male brains and three adult brains (Brodmann area 46) in the manuscript were similar, indicating reproducibility. While average barcode methylation remained stable with age for inhibitory neurons, upper and lower cortical excitatory neurons exhibited significantly higher methylation levels in older brains. This trend suggests a preferential loss of barcode methylation in “older” excitatory neurons, as significantly fewer neurons with less methylated barcodes were sampled from older brains (Fig 6C &6D). Alternatively, if fCpG methylation remains stable after cell division ceases, the observed decrease in average barcode methylation could indicate that neurons with less methylated barcodes are preferentially lost with aging. This suggests that neurons arising later in development may have higher mortality during aging. While other explanations are possible, the data illustrate how barcodes could be potentially used to infer both brain development and aging. Overall, these two new datasets help illustrate the degree of fCpG barcode reproducibility, and stability during aging.

**Summary**

This supplement addresses several concerns raised during the Public Review. The criteria for the fCpG barcode, as stated in the manuscript, are as follows: 1) a defined initial pattern in a progenitor cell; 2) polymorphic changes upon cell division; 3) sufficient polymorphism to distinguish between most cells; and 4) the capability to record ancestry. Among these criteria, only the adequacy of barcode polymorphisms to differentiate between most inhibitory and excitatory neurons has been well-established, indicating that while evidence for lineage tracing is supportive, it remains inadequate and dependent on assumptions that are difficult to validate.

The supplement provides the requested more detailed explanation of barcode mechanics. Simulations that begin with a single cell containing a fully methylated barcode broadly align with experimental data, incorporating relatively straightforward cell divisions, deaths, and expansions alongside random barcode replication errors. The brain facilitates serial barcode examinations because neurons typically stop dividing at different times during development. New barcode data reveal that daughter cells exhibit greater relatedness during embryogenesis, even amidst active and passive demethylation. Additionally, new data demonstrate the degree of barcode reproducibility and stability throughout aging.

Tools to reconstruct human brain development are limited, and fCpG barcodes can potentially add complementary information such as mitotic ages, diversity within a clade, and migration patterns between immediate daughter cells. fCpG barcode complexity is required because most brain development occurs prenatally, necessitating very high replication error rates to distinguish between billions of adult cells. Such high error rates (~$10^{-2}$) would inherently lead to polymorphic methylation patterns, complicating the differentiation of ancestral signals from technical or biological noise, especially if methylation remodeling occurs independently of cell division. While the current evidence for fCpG barcode lineage tracing is inadequate, a functional DNA methylation-based barcode comprising as few as 50 fCpG sites (yielding $10^{15}$

unique binary patterns) with similar high replication error rates could feasibly reconstruct aspects of human brain development and aging.

#### New Supplement Methods

**fCpG barcodes during embryogenesis:** The data are from GSE100272 (2). All oocyte data were used. Male zygotes and embryos were identified by greater numbers of Y chromosome reads. Average fCpG coverage was 2,268 reads per cell with a minimum of 897 reads. PWD comparisons required at least 25 comparable fCpG sites between cell pairs. The subset of cells used is provided in new Supplemental File 1.

**fCpG during aging:** The data and cell annotations were from refs 5 and 6, with cells used for analysis in the new Supplemental File 1. There were at least 400 fCpG sites per cell, and at least 25 comparable fCpG sites between cell pairs for PWD calculations.

**Pipelines**
(Files are in Pileline.zip)
1: Download GSE file from GEO
2: Filter out smaller files (typically keep 70 mB or larger tsv.gz files)
3: Use Program 1 (“mergetsvgzcol123divide4by5.py” with “locALLforfirstfilter.csv”) to select CpG sites at the chromosome locations (+ and - strands) of the ~31k fCpGs, and calculate methylation. A .tsv.gz file produces a .tsv file.
4: Use Program 2 (“collectmasterlistfromsmalltsvfastermerge” with “master_list.csv) to make a csv file with the methylation of each fCpG site for each cell, with blank sites labeled as “NA”
5: Use Excel to annotate single cells with their phenotypes from the published keys (1,5). Also calculate the fCpG methylation, and number of fCpGs with data for each cell.
6: Filter cells without relevant phenotypes, and cells with insufficient numbers of fCpG sites with data (500 for the manuscript and 400 fCpG sites for the new Supplement).
7: Calculate PWDs and the number of fCpGs that are compared between all possible cell pairs. Program 3 (“pwd2.py”) compares between all cells in a single .csv file (no headers, with individual cells across and fCpGs down). Program 4 (“pwdsquare.py”) calculates PWDs and the number of fCpGs compared between cells when two .csv files are compared. Programs 3 and 4 were used to segment the data into smaller chunks to shorten run times. Each program produces a csv file with PWDs and a csv file with numbers of fCpGs compared.
8: Excel was used to remove cell pairs with too few comparable fCpGs (minimum of 30 for the manuscript and 25 for the new Supplement).
9: Excel was used to calculate average PWDs between all comparable cell pairs and to find the minimum PWD of its nearest neighbor.

**Simulations of Neurogenesis**
Program 5 (“NumPydet_lin_fCpG_5_1.3arrayflipavemultrun_oneout_good_writeN_XeditDS.py”) is used to simulate different growth scenarios and is parameterized with two csv files. The first csv file (“lookup5_table_Xonly.csv” with four columns; lineage, array1, p0_1, p1_0) initializes the first progenitor cell. The number of rows is the number of fCpG sites, array1 is the starting methylation state (all 1’s), p0_1 is the probability of flipping from 0 to 1, and p1_0 is the probability of flipping from 1 to 0 and is 0.01 in the simulations. The other csv file (“cellnumber_table.csv” with five columns; division, desired_population, q, r, s) controls cell population size and cell death with each division. The desired_population provides the number of cells after each cell division. The program determines cell survival by randomly selecting (without replacement) for each mother cell a q, r or s value, where (q+r+s) = (the number of mother cells). A cell with “q” will have one surviving daughter cells, “r” has 2 surviving daughter cells and “s” has no surviving daughter cells.

The outputs are “sumlineage_arrays.csv” that calculates the average methylation of the cell population at each fCpG site, and “last_run_lineage_arrays.csv” which outputs the methylation of each fCpG site for each simulated cell. Program 6 (“covertrowlosecommas.py”) produces “out.csv” to format the data for Program 7 (“pwdofNcellsNaN4sigAVE_Rruns.py”) that samples specific numbers of simulated cells to match the numbers of cells sampled with the experimental data. Program 8 (flip101010.py) was used for the simulations of Fig 2.

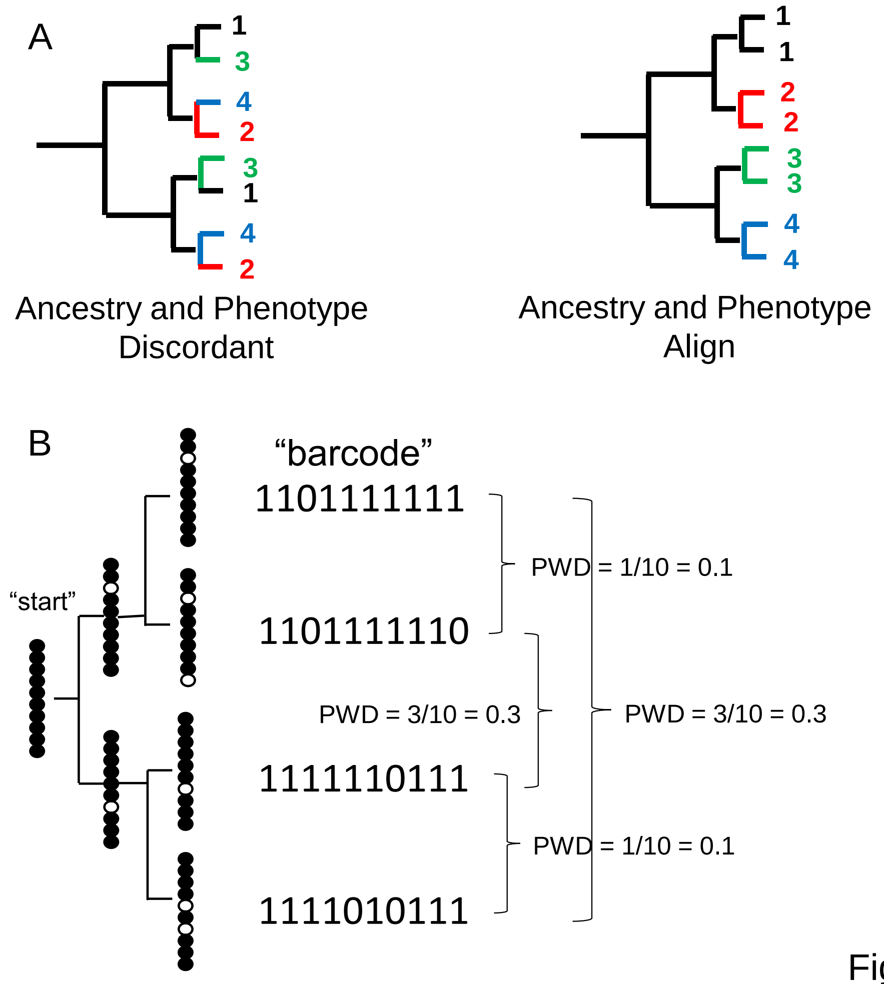


Figure 1

**Figure 1) A:** Trees can be reconstructed by comparing phenotypes or by comparing genomic differences such as fCpG barcodes. Ancestry and phenotypes may be discordant if progenitor cells produce cells of different phenotypes. More typically, ancestry and phenotypes align because cells with the same phenotypes tend to have common progenitors. For the single cell brain data, ancestry and phenotype align because cells of the same type are generally more closely related.

**B:** fCpG barcodes appear to start predominately methylated in the progenitor cell. With division, random replication error occur and are propagated to daughter cells. Counting and then averaging the differences between fCpG sites yields an average pairwise distance (PWD, range 0 to 1). More related daughter cells tend to have lower PWDs, but barcodes may also match by chance.

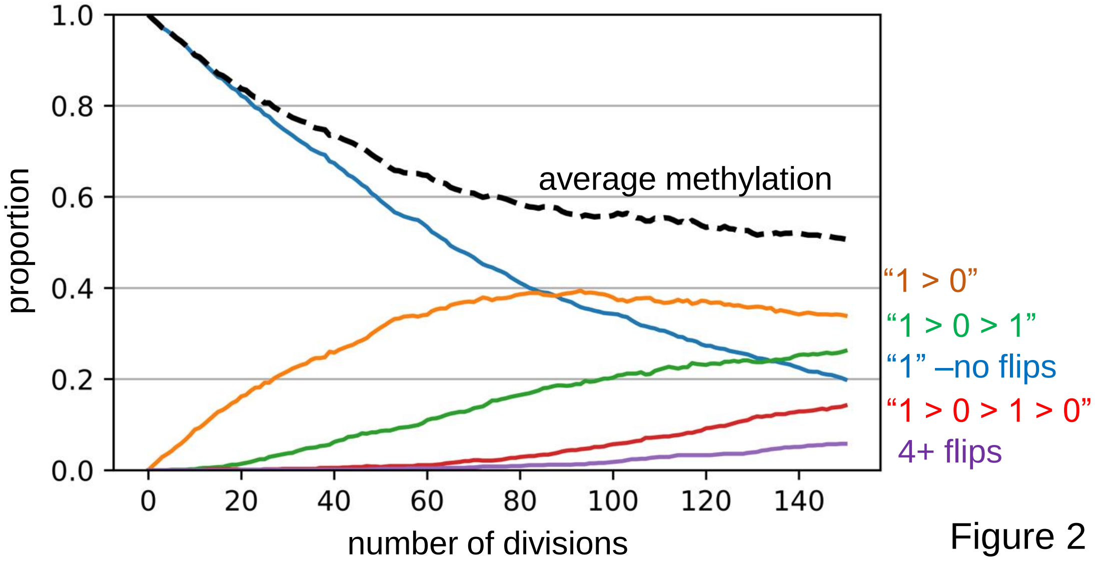


Figure 2

**Figure 2)** Barcode dynamics. Simulations broadly consistent with the experimental data indicate a replication error rate of 0.01 per fCpG site per divsion, with equal probabilities of changes or flips from methylated to demethylated (1 > 0) and from 0 > 1. A simulation for excitatory neurogenesis is shown, where simplistically, excitatory neurons cease division and appear after 150 divisions. The graph displays how individual fCpG sites change through time. The fCpG barcode starts methylated, and barcode methylation decreases with divisions. Even with an error rate of 0.01, after 150 divisions only about 5% of fCpG sites experience four or more flips, and half have had zero or only a single flip. A fCpG barcode can still effectively distinguish between cells if the flips are random and multiple fCpG sites are compared between cells. Although backflips complicate analysis, the large numbers of replication errors facilitate comparisons between neurons that develop during a short prenatal interval.

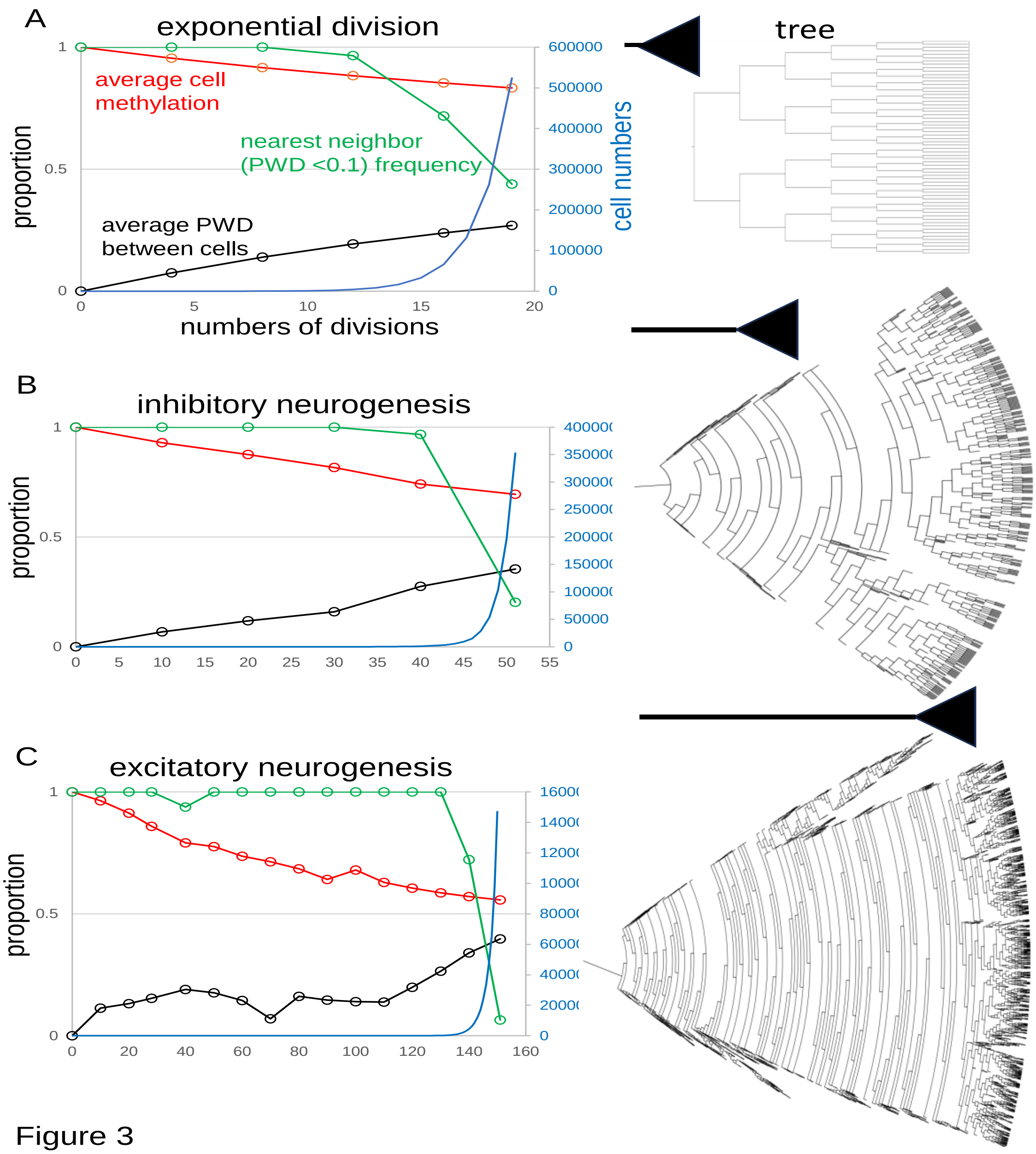


Figure 3

**Figure 3)** fCpG barcode simulations. The simulations start with a single progenitor and a fully methylated barcode with 200 fCpG sites and an error rate of 0.01 per division. At each time point, up to 1,000 cells are sampled from the population to calculate fCpG barcode average methylation, average PWD, and cell proportions with a nearest neighbor with a PWD <0.1. Final tree expansions are truncated to allow for visualization. A range of simulations are broadly consistent with the experimental data.

**A:** Simulation of exponential growth where each cell yields two daughter cells broadly models early hindbrain neurogenesis. After 19 divisions, average barcode methylation is ~0.83, PWD is ~0.27, and among the 1,000 sampled cells, for about 44% of cells there is another cell with a similar barcode (PWD <0.1).

**B:** Simulation of inhibitory neurogenesis with differentiation after 50 divisions. Early divisions are characterized by cell death (zero or one daughter, represented by dead ends in the tree), with terminal growth. After 50 divisions, average barcode methylation is ~0.69, average PWD is 0.35, and 20% of sampled cells have a nearest neighbor (PWD <0.1)

**C:** Simulations of excitatory neurogenesis with differentiation after 150 divisions. As with inhibitory neurogenesis, cell death during early divisions limits population size before terminal expansion. After 150 divisions, average methylation is ~0.56, PWD is ~0.4, and ~6% of cells have a nearest neighbor (PWD <0.1) among the 1,000 sampled cells.

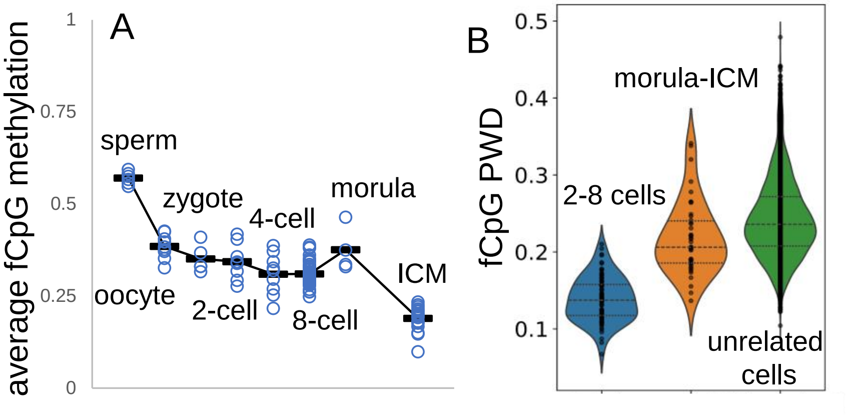


Figure 4

**Figure 4)** fCpG barcodes at conception, when germline methylation is erased by active and passive demethylation. Whole genome bisulfite single cell sequencing data are from GSE100272 (4). Male cells were inferred from a paucity of Y chromosome reads.
**A:** fCpG methylation generally decreases during early development. fCpG barcode methylation is highest in sperm, albeit sperm X chromosomes yield female zygotes. A brain cell progenitor with predominately methylated fCpGs was not evident.
**B:** Unlike at the start of brain development, fCpG barcodes early in life are polymorphic between unrelated embryos. However, fCpG barcodes are more similar between related cells in 2 to 8 cell embryos, and less similar between cells in morulae and the ICM. Dots indicate values of cell pairs, with a minimum of 25 comparable fCpG sites.

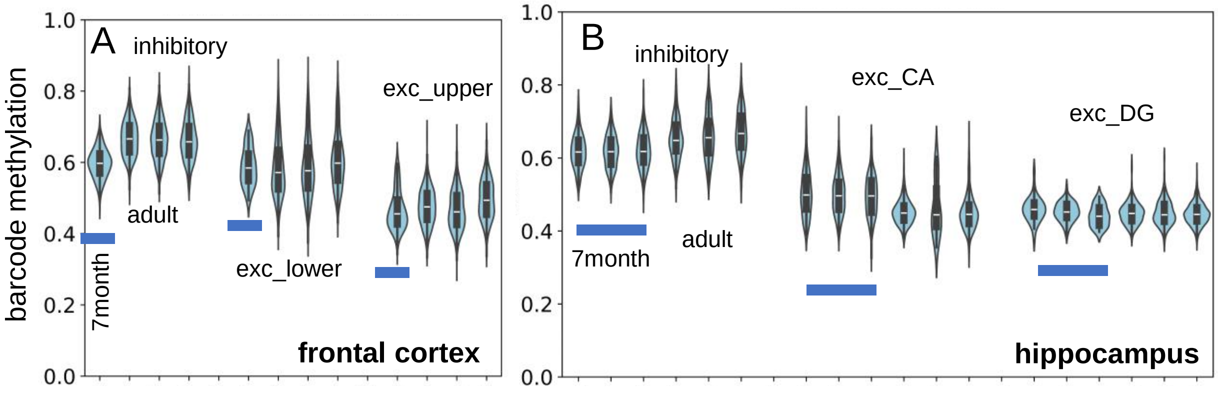


Figure 5

**Figure 5)** New single cell data (5) indicate fCpG barcode methylation at 7 months of age is similar to adult levels (H02, 29 yo; H01, 42 yo; H04, 58 yo).
**A:** Inhibitory, and lower and upper cortical excitatory neuron barcode methylation levels from the frontal cortex are similar.
**B:** Inhibitory and excitatory (CA and DG) neuron barcode methylations levels are similar between infant (3 samples) and adult hippocampus.

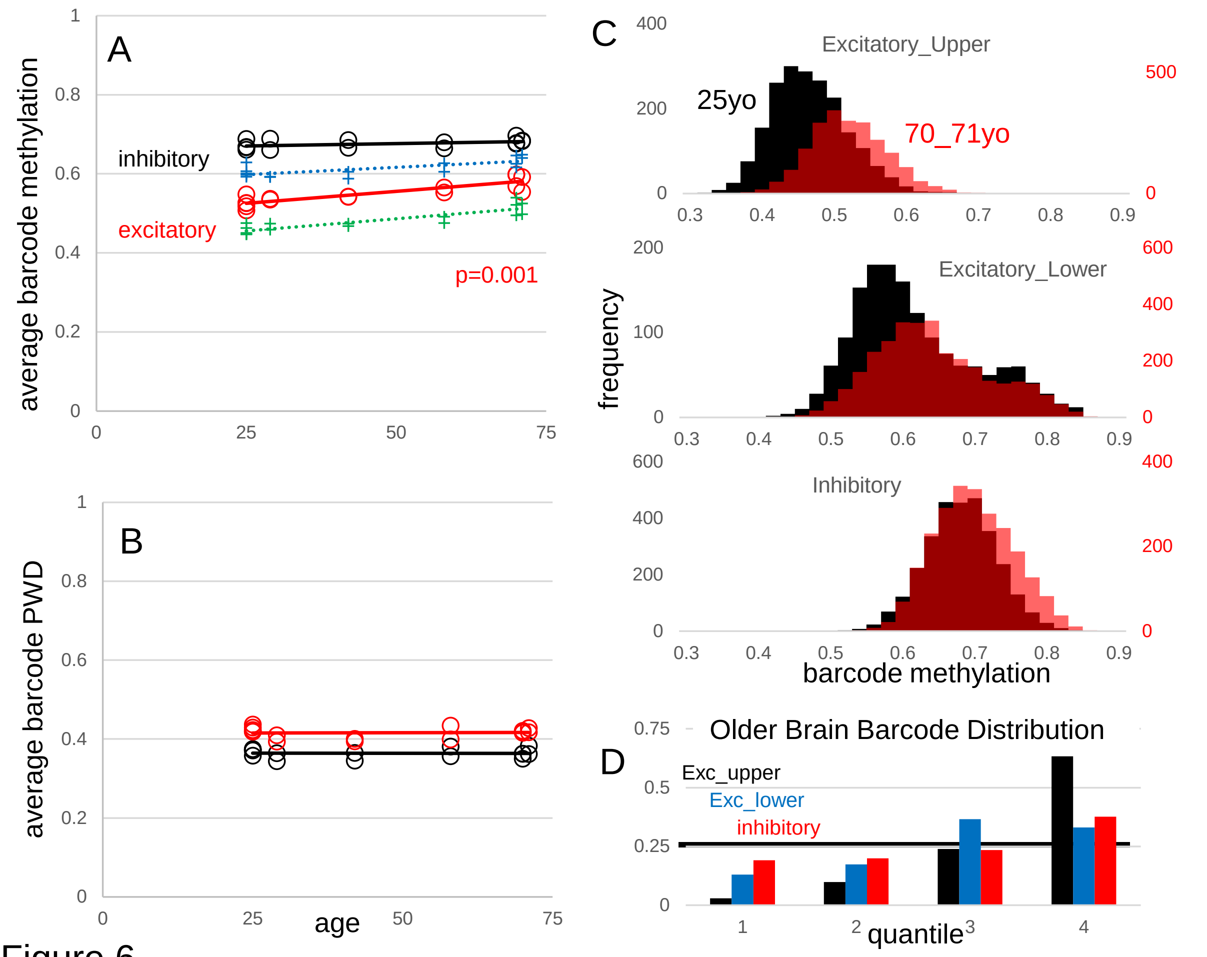


### Figure 6

**Figure 6)** fCpG barcode reproducibility and stability with aging. New data (6) are frontal cortex (Broadman area 46) WGBS single cells from young (25 year-old, 3 individuals and 5 samples) and older (70-71 year-old, 3 individuals and 5 samples) males. Data from the manuscript are also shown (H02, 29 yo; H01, 42 yo; H04, 58 yo).

**A:** Average neuron barcode methylation levels were similar between different aged individuals. Excitatory neuron barcode methylation in older males (70-71 yo) was significantly greater than the 25 yo males (t-test comparing only the 25 yo and 70-71 yo groups). Both upper (green) and lower (blue) cortical excitatory neurons showed greater average barcode methylation with aging.

**B:** Average inhibitory and excitatory neuron barcode PWDs (comparing within subtypes in each brain) were similar between different aged individuals, indicating that barcodes remain polymorphic.

**C:** Composite histograms of individual neuron barcode methylation levels for younger (black, 25yo) and older (red, 70-71yo) brains. There is a preferential loss of neurons with less methylated barcodes, especially with excitatory neurons .

D: Younger brain neuron barcodes were used to define quantiles. Older brain neurons with less methylated barcodes were depleted in the less methylated quantiles, with significant differences for all quantiles (Mann-Whitney U test, $p < 10^{-9}$).
